# The mutational landscape of innervation axes in cancer

**DOI:** 10.64898/2026.07.21.739760

**Authors:** A. Batool, C. Arora, L.A. Nemati Fard, R. Passannanti, M. Varisco, R. Vukotic, S. Capsoni, A. Cattaneo, F. Raimondi

## Abstract

The mutational landscape of neural innervation axes in cancer has garnered increasing attention due to their significant influence on tumorigenesis, metastasis, and treatment resistance. This study provides an overview of the alterations of neural signaling pathways based on multi-omics data analysis. In particular, we conducted a comprehensive analysis of mutations in key genes associated with neurotrophin signaling, norepinephrine and cholinergic pathways, alongside paracrine and synaptogenic factors. Our findings reveal that 65% of patients in a pancancer cohort exhibit at least one somatic alteration in these pathways, with neurotrophin genes being the most frequently altered. Notably, alterations in these genes (e.g., NGF) correlate with poor patient survival across various cancer types, suggesting their oncogenic potential. We also identified significant co-occurrence of mutations and differential expression patterns from both bulk and single-cell RNAseq data, indicating complex interactions among these pathways that contribute to tumor innervation and neuroplasticity.

This study underscores the necessity of understanding the crosstalk between neural signaling and cancer biology to improve patient stratification based on mutational profiles and identify potential therapeutic targets. Further research utilizing advanced genomic techniques is essential to elucidate the mechanistic roles of these pathways in tumor innervation.

## Introduction

The role of neural innervation axes in cancer is emerging as a new hallmark of cancer(1). Recent investigations have revealed that neural signaling pathways influence metastasis, tumorigenesis, and treatment resistance (2). The function of somatic mutations in the neural pathway, particularly related to neurotrophic factors, neurotransmitters, and synaptic plasticity, has provided insight into the molecular and genetic characterization of innervation axes. Despite advancements in the research of the individual signaling pathways, the interplay of these pathways remains inadequately characterized, thus preventing synergistic, precision therapies.

Prior research has shown that cancer innervation is influenced by a feed-forward loop involving cancer cells, neurons, and stromal factors. For example, NTRK neurotrophin receptors have been recognized as oncogenic drivers, whereas the loss of function of synaptogenic factors such as *THBS1* seems to hinder inhibitory synaptic signals, thereby promoting hyperinnervation (3). In addition, paracrine signaling molecules such as *COL1A2* (collagen) (4) and *NLGN3* (neuroligin-3) (5) modify the extracellular matrix in ways that facilitate axon guidance. Mutations in other key regulators such as *PTCH1*, a modulator of the Hedgehog pathway, and *GPC3*, a glypican proteoglycan can further disrupt synapse construction and alter the availability of growth factors (6,7).

Although individual gene alterations may initiate unique oncogenic processes, tumorigenesis is rarely driven by a single molecular event (8). Cancer rather emerges from the simultaneous disruption of hallmark processes and pathways, which together alter cellular activities, modulate responses to microenvironmental stress, and complicate therapeutic interventions (9). In the context of tumor innervation, some studies have begun to reveal the crosstalk between neuronal signaling pathways. Notably, a bidirectional feed-forward loop involving neurotrophin signaling with norepinephrine (*NGF-ADRB2/3*) (10) and acetylcholine (*NGF-CHRM1/3*) (11) has been implicated in inducing neuronal infiltration and perineural invasion in pancreatic and gastric cancers. These landmark studies opened new avenues for synergistic targeting of innervation pathways in cancer.

This study provides a landscape analysis of the mutations of genes participating to pathways established in tumor innervation, including neurotrophins, norepinephrine and cholinergic signaling pathways, which collectively regulate neuronal survival, axonal guidance, and neurotransmitter release, which are processes co-opted by tumors to innervate and cause neuroplasticity. Additionally, we investigated mutations in paracrine neuronal factors and synaptogenic factors, which influence the structure and functional adaptation of nerves within the tumor microenvironment (2). We systematically investigated mutations and signatures of innervation genes from cancer genomics datasets to further highlight novel mechanisms underlying tumor innervation as well as to uncover new biomarkers for personalized oncology. The approach is essential for understanding how tumors utilize neural plasticity, identifying combinatorial treatment targets, and forecasting resistance mechanisms ultimately facilitating the disruption of the neuro-tumor symbiosis that underlies cancer proliferation.

## Results

### The mutational landscape of innervation genes

We curated a list of genes involved in tumor innervation and inspected the type and quantity of somatic mutations from cancer genomics datasets affecting them (see Methods). We considered the following groups of genes deemed as more important for tumor innervation based on literature review : neurotrophins (*NGF, NGFR, BDNF, IGF1, GDNF, ARTN, NTF3, NTF4, NTRK1, NTRK2, NTRK3*), norepinephrine signaling (*ADRB2, ADRB3, GNAS, DBH, TH*), acetylcholine signaling (*ACHE, CHRM1, CHRM3, CHAT*), paracrine neuronal factors (*COL1A2, NLGN3, NF1*) and synaptogenic factors (*GPC3, PTCH1, THBS1*).

We found a total of 1765 (60%) unique samples and 1676 (65%) unique patients from a pancancer cohort (i.e. (12)), carrying at least one somatic alteration, i.e. either simple mutations, copy number variations (either amplifications or homozygous deletions) and differential expression (DE; either significantly up- or down-regulations).

The most mutated genes are *NTRK1*, *CHRM3* and *COL1A2*, each one mutated in 14% of patients (Supplementary Figure 1). When considering individual pathways, the neurotrophin one is the most affected with 42% patients carrying at least one alteration, followed by genes mediating norepinephrine signaling (28%), paracrine neuronal factors (28%), acetylcholine signaling (26%) and synaptogenic factors (19%) (Figure 1A). We found a total of 78 significant instances of co-occurrence within the same sample of alterations affecting these innervation genes (FDR < 0.05). The most significant instances (Log2 Odds Ratio > 3, FDR < 0.001) are *ACHE - COL1A2, NTRK1 - CHRM3, GPC3 - NLGN3 and NTRK3 - NLGN3* (Supplementary Figure 1,2, Supplementary Table 1). On the other hand, we only found three significant instances of mutually exclusive alterations (FDR < 0.05): *NTRK2 - IGF1*, *NTRK3 - ADRB2* and *NTRK3 - TH* (Supplementary Figure 1,2; Supplementary Table 1).

**Figure 1:**
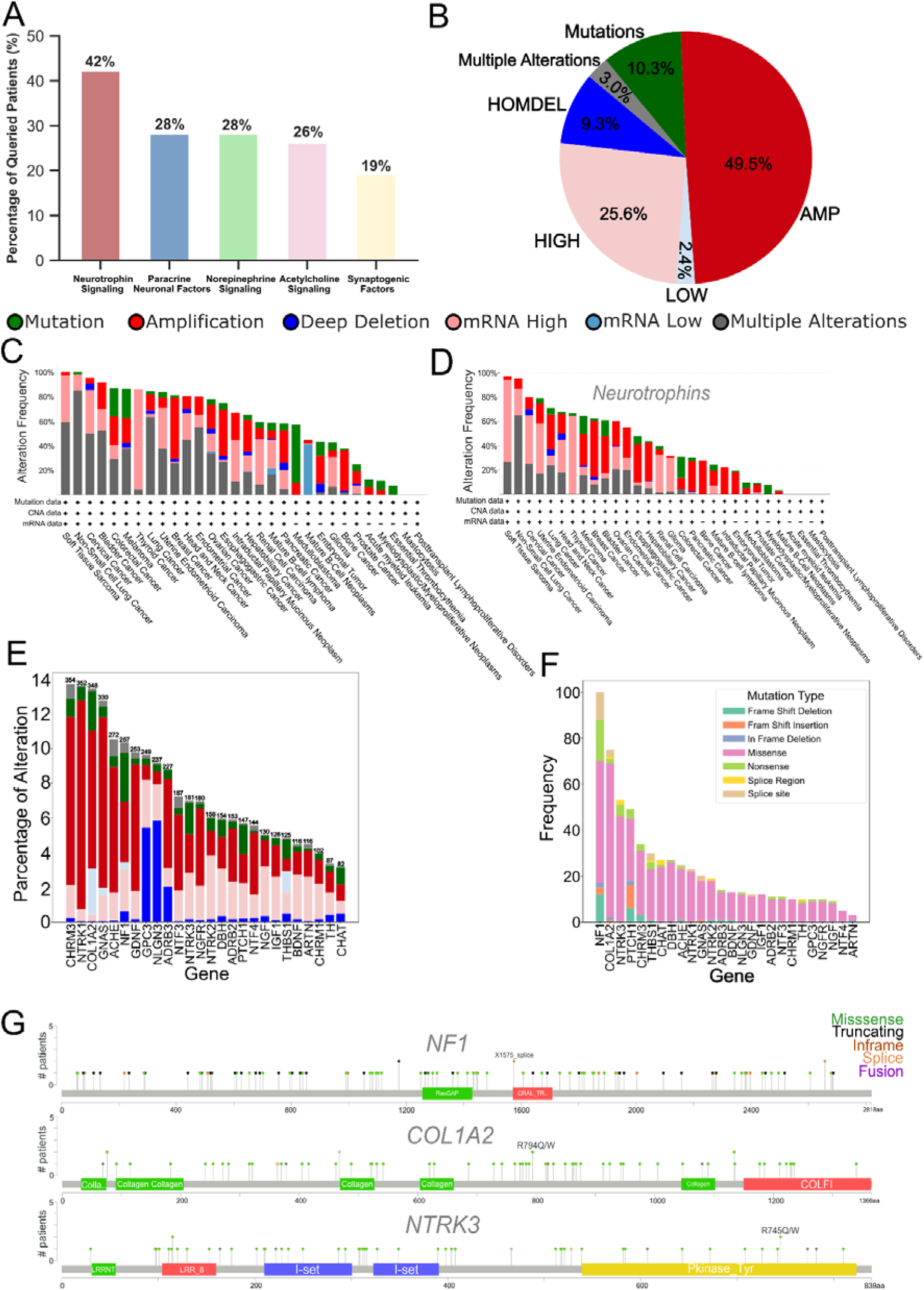
Summary of the distribution and frequency of alterations of innervation genes. A) Bar plot illustrating the percentage of patients with genomic alterations across the different innervation pathways. B) An overview of copy number variations (CNVs) and mRNA alterations across Pan-Cancer is reported. The data depicts the frequency of various alterations among all samples. C) The distribution of alterations in genes linked to innervation pathways across different types of cancer is illustrated. D) Distribution of genetic alteration in the neurotrophin pathway genes. E) Types of somatic mutations of genes in Pan-Cancers. The number represents the samples altered with the somatic mutations. F) Mutation frequency throughout pan-cancer for genes associated with the innervation pathway. The number shows the patients impacted by CNVs and mRNA alterations. G) A lollipop plot depicting mutations in particular genes.

We found a predominant prevalence of amplifications and higher expression of innervation genes, accounting for roughly 75% of total alterations, while Homozygous deletion and low expressions account for 13% and simple mutations for 10% of total alterations (Figure 1B). Considering altogether the genes participating in innervation pathways, the most affected cancer types are soft tissue sarcomas and non-small cell lung adenocarcinomas, with nearly the totality of samples carrying at least one alteration (Figure 1C). Blood tumors such as Acute Myeloid Leukemia, Myeloproliferative Neoplasms and Essential Thrombocythemia have little mutational burden for these genes. While the majority of tumors showed a major contribution of amplification or higher expression of innervation pathway genes, specific tumors showed prevalence of certain types of alterations, such as simple mutations in Medulloblastoma or low expression in Mature B-cell neoplasms (Figure 1C). Certain pathways, such as neurotrophins, display a prevalence of amplifications or high expressions in every cancer type considered, suggesting a potentially universal oncogenic role (Figure 1D). The most altered gene in the neurotrophin pathway, i.e. *NTRK1*, is consistently amplified across tumors and overall altered in 13% of considered samples (Figure 1E), and it’s mutated in nearly 30% of the samples of Breast carcinoma, Soft Tissue Sarcoma, Non-Small Cell Lung Cancer Hepatobiliary Cancer cancer (Supplementary Figure_3). Others, such as synaptogenic factors, display instead a prevalence of homozygous deletions (e.g. *GPC3* and *NLGN3*) (Figure 1E, Supplementary Figure 4), suggesting a potential tumor-suppressive role.

When considering simple mutations, *NF1* is the most mutated gene (Figure 1F), with mostly loss-of-function mutations distributed across gene’s architecture (Figure 1). The other two most mutated genes are *COL1A2* and *NTRK3* (Figure 1F,G).

We considered gene fusions from a compendium of genetic rearrangements (i.e. FusionGDB2.0), which revealed recurrent structural alterations across a variety of cancer types, as evidenced by the number of unique patient samples affected by fusions (Figure 2A). *GNAS* was identified as the most frequently fused gene, occurring in more than 130 unique samples, indicating its potential role as a locus for genomic instability. Other genes exhibiting elevated fusion counts include *NF1, COL1A2, NTRK3, NTRK1*, and *THBS1*, each associated with multiple distinct fusion partners across diverse tumour types.

**Figure 2:**
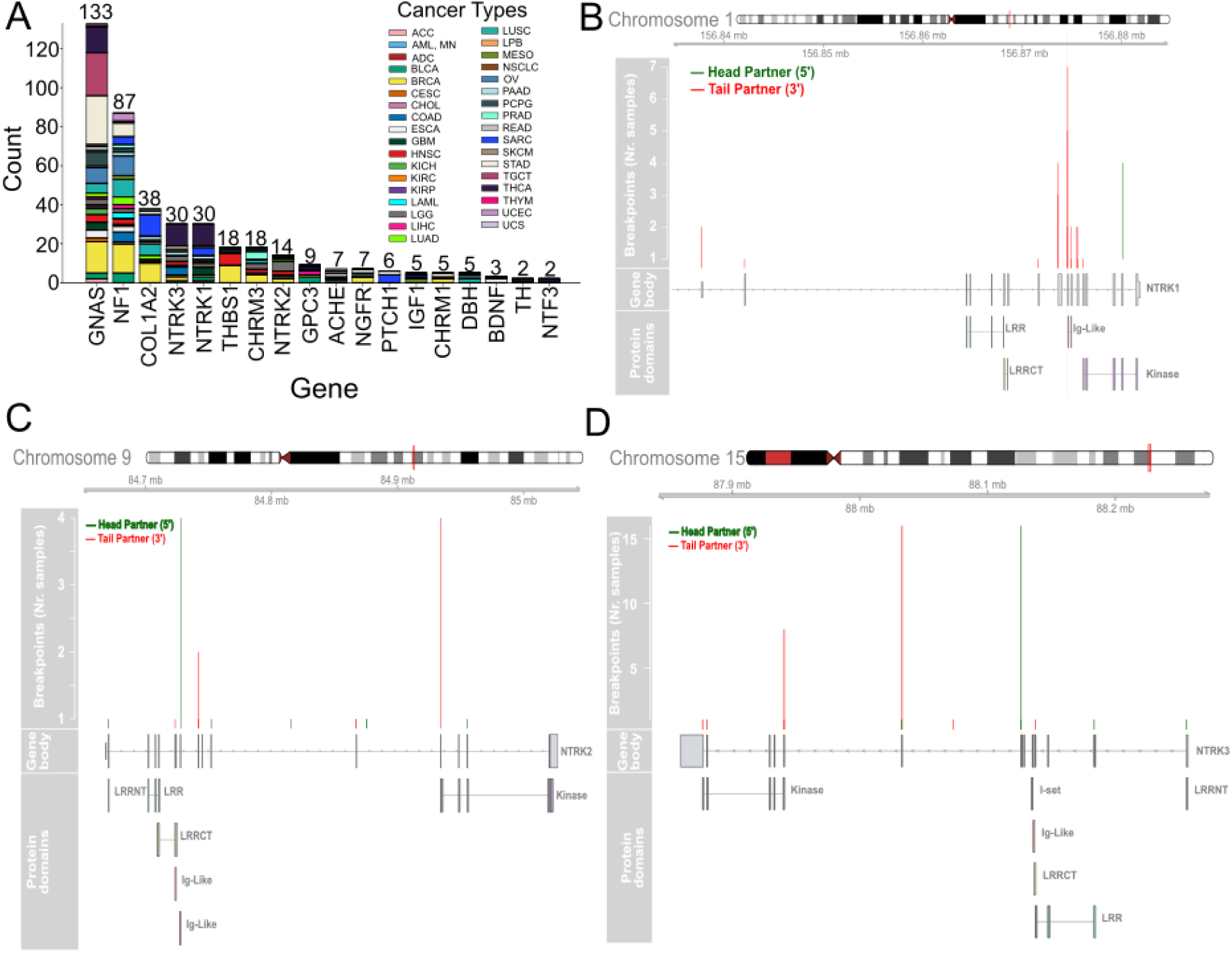
Pan-cancer fusion landscapes and genomic breakpoint profiles of the NTRK gene family. A stacked bar chart illustrating the distribution of unique sample counts containing gene fusion events across several target genes, categorized by cancer type (retrieved from FusionGDB2.0). B) Genomic representation of the positive-strand NTRK1 locus, illustrating 5′ (head, green) and 3′ (tail, red) breakpoint sample counts in conjunction with Ensembl transcript architecture. Red vertical bars emphasize concentrated breakpoint hotspots. C) Genomic mapping of the NTRK2 locus, depicting the sample counts of associated breakpoints and the limits of structural genes. D) Genomic mapping of the minus-strand NTRK3 gene, with structural track coordinates reversed to appropriately represent the negative strand orientation and transcriptional direction. All genomic locations and transcript mappings are derived from the GRCh38 assembly. In all genomic tracks (B–D), Pfam protein domains are consistently color-coded as outlined below: Leucine-rich repeats (LRR) depicted in green, LRR C-terminal (LRRCT) illustrated in yellow, Immunoglobulin-like (Ig-like) represented in salmon, and the Tyrosine Kinase domain shown in purple.

Neurotrophin receptors *NTRK3* and *NTRK1* were identified as the most frequently affected by fusions in the neurotrophin pathways, consistent with their recognised function as oncogenic drivers in multiple tumour types. The fusions create constitutively active TRK proteins that facilitate tumorigenesis and have been identified in thyroid carcinoma, gliomas, and sarcomas, functioning as diagnostic biomarkers and therapeutic targets (13). Consistent with these observations, *NGFR* was identified as a fusion partner, though at a lower frequency, indicating a more context-specific role. Ligand genes, including *BDNF* and *NTF3*, were less frequently affected by fusions in a tumor-specific fashion (Figure 2A).

We systematically characterized the structural landscape of these oncogenic rearrangements by mapping the distribution and orientation of genomic breakpoints on the domain architectures of NTRK1, *NTRK2* and *NTRK3*. Structural analysis revealed significant locus-specific preferences and distinct fusion partners for the three receptor tyrosine kinase genes, aligning with classical models of TRK kinase activation. *NTRK1* structural rearrangements demonstrated a dense clustering of breakpoints localized within the extracellular Immunoglobulin-like (Ig-like) domains and the catalytic Tyrosine Kinase domain, largely acting as 3′ tail partners. This configuration ensures the maintenance of the intracellular kinase domain, enabling ligand-independent dimerisation and continuous downstream signalling when fused to an upstream 5′ head driver (Figure 2B). This pattern is seen in the recurring *TPM3-NTRK1* fusion reported in many malignancies (13), consistent with pan-cancer fusion profiling of kinase genes (14).

Conversely, the breakpoints of *NTRK2* are more scattered, with a major hotspot of susceptibility in the amino-terminus of the extracellular domain, specifically in the Leucine-Rich Repeat N-terminal (LRRNT) and Leucine-Rich Repeat (LRR) regions, and mostly acting as a 5′ head partner (Figure 2C), as observed in the *STRN-NTRK2* fusion identified in soft-tissue sarcoma (15).

NTRK3 exhibited breakpoints mapping exclusively to the catalytic Tyrosine Kinase domain as 3′ tail partners, a conformation essential for producing the classic constitutively active oncoprotein driven by upstream dimerisation anchors, such as the well reported *ETV6-NTRK3* driver (16). In contrast, the breaks of its 5′ head partner were strictly confined to the upstream regulatory machinery, encompassing the LRRNT, Leucine-Rich Repeat C-terminal (LRRCT), LRR, and Ig-like domains (Figure 2D).

### Tumor innervation pathways expression signatures

We checked the differential expression (DE) profile of innervation genes by comparing cancer tissues (from TCGA) against matched normal tissues (from GTEx; see Methods). We found that all the considered genes are differentially expressed in at least one tissue. The most up-regulated gene across tissues is *GNAS*, which is found over-expressed in 15 tissues out of 16 considered (supplementary Figure 5), while the tissue with the highest number of innervation genes differentially expressed is pancreas (supplementary Figure 5). *NTRK3* is down-regulated in every tissue, consistent with the established role of the *NTF3-NTRK3* axis as a tumor suppressor in several cancer types (17) (supplementary Figure 5). In pancreas, we found up-regulated many genes of the recently described feedforward loop between neurotrophin and Beta2 Adrenergic receptor-PKA pathways (10), including *NGF* and *BDNF* neurotrophins, beta adrenergic receptors *ADRB2*, *ADRB3*, the Tyrosine 3-monooxygenase *TH*, the bottle neck enzyme for catecholamine biosynthesis, in addition to the Galpha s subunit *GNAS* (supplementary Figure 5). Other tissues with up-regulation of innervation genes are Testis, Lung and Uterus.

We also explored pre-calculated differential expression (DE) from a compendium of single-cell RNA-seq datasets from multiple cancer types (i.e. Tisch2 (18), see Methods), which revealed that the neurotrophin pathway genes were differentially expressed in most of the tissues. We found that *NTRK2* and *NGFR* were the most dysregulated genes across a wide range of tissues and cell types (Figure 3A). In particular, *NTRK2* and, to a less extent, *NGFR* and *NTRK3*, were found down-regulated across multiple tissues. *NTRK3* and *NTF3* were down-regulated in the brain, whereas *BDNF* exhibited predominant down-regulation in the skin (Figure 3A). On the other hand, *NTRK1* and *NGF* are widely up-regulated across cancer tissues, with the only exception of down-regulation of *NTRK1* in certain pancreatic cancer cells (Figure 3A,B).

**Figure 3:**
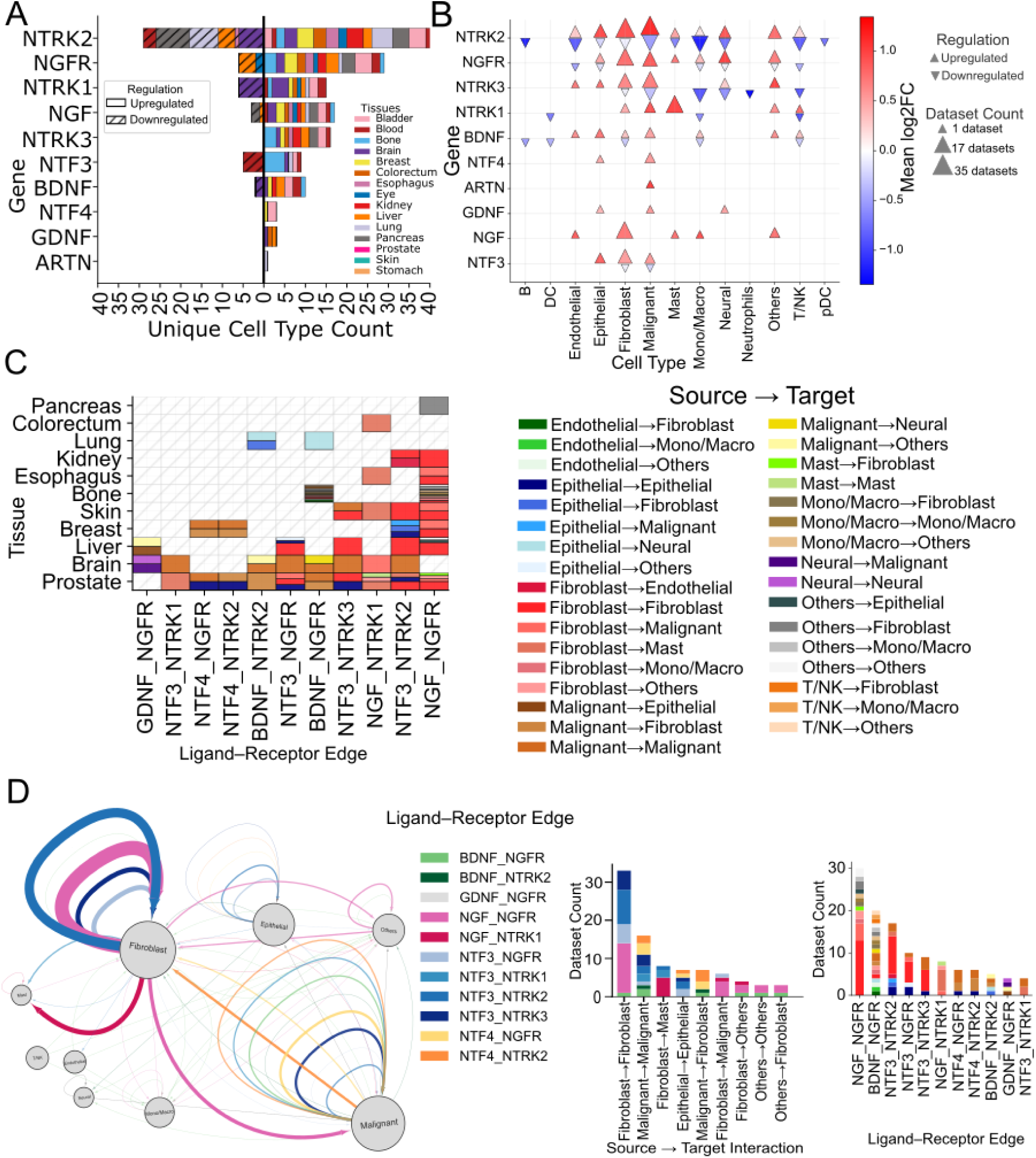
Differential expression of neurotrophin pathway genes in the Tisch single-cell RNA-seq database. A) The stacked bar plot depicts the count of unique major cell type categories of neurotropic genes across various tissues. The up-regulated gene is shown as a solid colour bar, whereas the down-regulated gene is illustrated as a hatched colour bar. B) The triangular scatter plot illustrates the distinct dataset counts and mean log fold change for each gene across various major cell types. Up-regulated genes are represented by upward-pointing triangles, while down-regulated genes are indicated by downward-pointing triangles. The size of the triangle indicates the number of unique datasets, while the colour intensity reflects the mean log2 fold change, with red signifying up-regulation and blue indicating down-regulation. C) The stack matrix plot shows the up-regulated ligand_recptor interactome in cell types of various tissues. The colored stacks indicate the particular source and target cell types for each interaction. D) The global signaling network including all tissues. The size of nodes and the size of labels are proportionate to out-degree to emphasize key signaling hubs, while edge thickness and transparency correspond to the number of datasets to signify interaction robustness along with a stacked bar plot depicting the distribution of source and target cell-type interactions within the relevant tissue.

We inspected the DE profile of each neurotrophin gene by aggregating DE analysis of every study for each tissue (see Methods). Cell types such as B cells, DC, pDC and Neurotrophils have little neurotrophin axes expressed, and the few ones are down-regulated (Figure 3B). Malignant cells or Fibroblast have a prevalence of up-regulated neurotrophin axes. *NTRK1* is consistently up-regulated in Fibroblast, Malignant, Mast cells, mixed up- and down-regulated in T/NK cells, and down-regulated in DC and Macrophages. *NTRK2* shows mix up- and down-regulation in most of the cell types, while *NTRK3* exhibits a selective profile with mostly up-regulation in endothelial, epithelial and stromal cells, mixed up- and down-regulation patterns in malignant and neural cells, and mostly down-regulation in macrophages, neutrophils, and T/NK cells (Figure 3B).

Among the ligands, *NGF, NTF4*, *ARTN* and *GDNF* were exclusively up-regulated (Figure 3B), though with distinct patterns: *ARTN* expression was highly specific to malignant cells (Figure 3B), particularly in lung cancer (Supplementary Figure 6), consistent with its role in RET-dependent tumorigenesis and metastasis (19), while *NGF* overexpression was mainly observed in fibroblast cells and, to a lower extent, in endothelial cells, mast cells and in monocytes/macrophages (Figure 3B). *BDNF* and *NTF3* display alternate patterns of cell type-specific up- and down-regulation. In particular, *BDNF* is down-regulated in B and DC cells (Figure 3B).

We considered functional interactions between neurotrophin and neurotrophin receptors available from the String Database. Most widespread interactions across cell types and tissues include *NGF_NGFR*, *NTF3_NTRK2*, and *NGF_NTRK1* (Figure 3C). *NGF_NGFR* signaling was predominantly mediated through fibroblast-to-fibroblast and fibroblast-to-malignant interaction. Certain tissues, such as the prostate, brain or liver, are characterized by several neurotrophin axes between different cell types (Figure 3C). Other tissues are characterized by specific neurotrophin axes, i.e. *NGF_NGFR* involving fibroblast cells in pancreas, *NGF_NTRK1* between fibroblast and malignant cells in colon and *BDNF_NTRK2* and *BDNF_NGFR* involving epithelial and neuronal (or epithelial and fibroblast) cells in lung (Figure 3C). *GDNF_NGFR* was only detected in the brain and liver tissue at neuronal-to-neuronal and neuronal-to-malignant interfaces in the brain, corresponding to GDNF established role in neuronal survival and glioma progression (20). Furthermore, *NTF4_NTRK2* and *NTF4_NGFR*, confined to prostate and breast tissues, exhibited parallel patterns defined by malignant-to-fibroblast, malignant-to-malignant, and epithelial-to-epithelial signaling (Figure 3C).

Overall, the most recurrent cell-cell communications mediated by neurotrophins are autocrine axes mediated by fibroblast and, to a less extent, by malignant cells. Communication between fibroblast and mast cells is the third most recurrent interaction (Figure 3D). The interactions between various neurotrophins (i.e. *NGF*, *BDNF* and *NTF3*) and *NGFR* are among the most recurrent cell-cell interactions (Figure 3D).

Overall, these findings imply that whereas individual neurotrophins (e.g., *NGF*, *ARTN*) are cell-type-specific in their expression, their receptors (*NTRK1*, *NTRK2*, *NTRK3*, *NGFR*) are heterogeneously but more widely regulated. The representation of all ligand–receptor pairs, e.g., *BDNF*–*NTRK2*, *NGF*–*NGFR*, *NGF*–*NTRK1*, and *NTF3*–*NTRK3*, highlights the existence of coordinated, cell-type–specific signalling loops which may support the notion of cell-type–specific autocrine and paracrine interactions in neuro-cancer communication.

As an example case study, we studied neurotrophin signaling networks in a scRNAseq atlas of Prostate adenocarcinoma (PRAD). In particular, we focused on the expression of *NGF* and its corresponding receptors throughout significant tumour microenvironment compartments as delineated by UMAP, encompassing malignant, epithelial, fibroblast, endothelial, and immune populations (Figure 4A).

**Figure 4:**
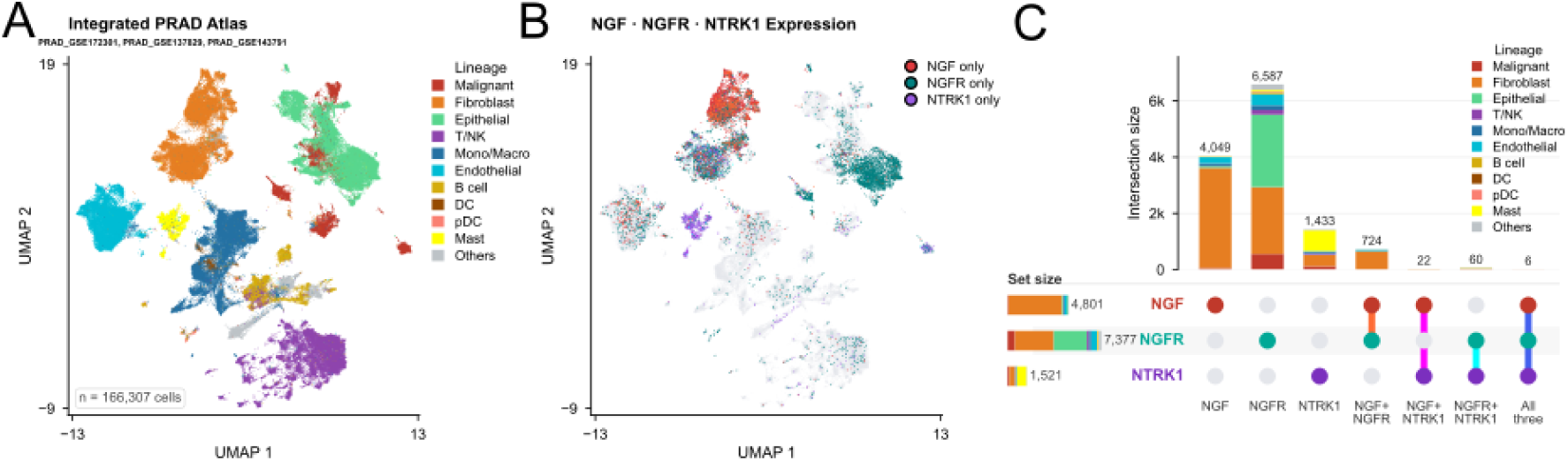
Single-cell expression profile and neurotrophin signaling network in the integrated PRAD atlas. A) UMAP embedding of the integrated PRAD single-cell atlas (n = 166,307 cells), batch-corrected using scVI and colored according to major cellular lineages. B) UMAP expression overlay emphasizing instances of single gene positivity (NGF, NGFR, and NTRK1) along the paracrine and autocrine axes. Gray points indicate cells that fall below the expression threshold (>0.1). C) Upset plot illustrating exclusive expression intersections (top) and matching total set sizes (bottom right) for each target gene, with bars stacked and colored by cell lineage.

The expression pattern of *NGF*, *NGFR* and *NTRK1* revealed a compartment-specific distribution. *NGF* (4,801 cells, 2.9% of cells) was mainly produced by fibroblasts, whereas *NGFR* (7,377 cells, 4.4%) was mainly enriched in epithelial and malignant cells. *NTRK1* (1,521 cells, 0.9%) was mainly located in mast cells (Figure 4B–C).

Co-expression analysis revealed two simultaneous signalling modalities. *NGF_NGFR* cells (n=724) were mostly fibroblasts (87.8%). In particular, it is possible to distinguish two fibroblast sub-populations, one enriched in NGF, and the other in NGFR expression (Figure 4B), suggestive of distinct signaling properties. *NGF_NTRK1* (n=22) and *NGFR_NTRK1* (n=60) cells were less frequent and more diverse, comprising fibroblasts, mast cells and malignant cells, indicating a limited paracrine pathway towards NTRK1 targets. Merely six cells exhibited co-expression of all three genes (Figure 4C).

These findings suggest that fibroblasts facilitate a dual-mode neurotrophin circuit, characterised by a predominant autocrine *NGF–NGFR* loop and restricted paracrine signalling to *NTRK1* expressing mast, malignant, and epithelial cells, hence supporting a stroma-anchored innervation-associated axis in this non-prostate tumour environment.

### Proteogenomics analysis of NGF and BDNF

In order to produce mature ligands, neurotrophins must undergo intracellular or extracellular proteolytic cleavage while disruption of this process can significantly change the signaling outputs between the survival and apoptotic pathways (21,22). We examined peptide distribution across functional domains to see if cancer tissues display different abundances of the propeptide (pro) and mature forms of NGF and BDNF could be detected (Figure 5A). In comparison to healthy controls, cancer cohorts are characterized by a better peptide coverage, especially across the Propeptide and Mature domains, according to linear peptide mapping (Figure 5A). BDNF and NGF mature regions produced a disproportionately higher number of distinct peptides in cancer vs healthy tissues (e.g., 65% for BDNF in cancer vs. 45% in healthy), indicating broader representation of mature-domain peptides in cancer proteomes which is likely connected to the higher number of proteogenomics studies for this condition (Figure 5B). Moreover, while in cancer tissues we observed a slightly higher number of distinct peptides mapped for the mature forms of neurotrophins, in healthy tissues we observed a higher number of unique peptides mapped to the propeptide forms (Figure 5B). Notably, these observations reflect differences in unique peptide diversity rather than peptide abundance and therefore complement, rather than replace, the quantitative PSM-based analyses described below.

**Figure 5:**
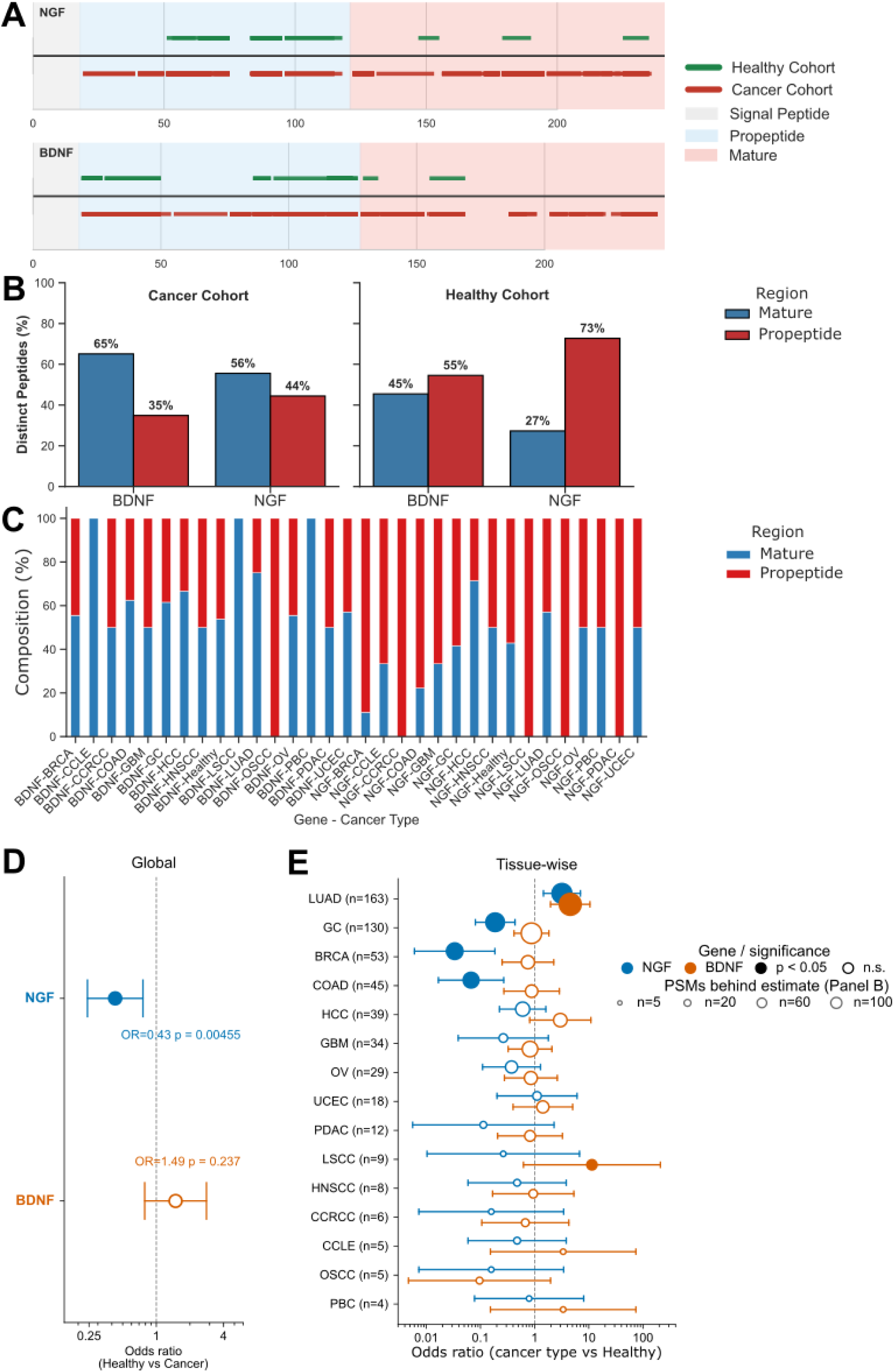
Comparative peptide mapping and domain distribution of NGF and BDNF in cancer and healthy cohorts. A) Linear sequence coverage plot shows identified peptides across functional domains (Signal Peptide, Pro-Peptide, and Mature) in cancer (red) and healthy (green)cohorts. B) The bar plots depict the percentage of unique peptides found within the Pro-Peptide and Mature areas of cancer and healthy datasets C) The bar plots show the relative composition (%) of mature vs propeptide fragments in various cancer types and healthy datasets. D) Comparison of pro-peptide and mature-domain peptide enrichment in NGF/BDNF using PSM based Fisher’s exact test, between healthy and cancer at a global level. E) Fisher’s exact test stratified by cancer types.

To formally test this compositional shift, we performed Fisher’s exact tests comparing pro-versus mature-domain peptide counts between healthy and cancer samples for each gene, using all confident peptide-spectrum matches (PSMs) (Figure 5D). Given the limited number of unique peptides identified for NGF and BDNF, PSM counts were used as the primary, adequately powered test. At the global level, NGF showed significant enrichment of pro-domain PSMs in cancer relative to healthy tissue (OR = 0.43, 95% CI 0.24–0.76, Fisher’s exact p = 0.0046), whereas BDNF showed no significant global difference (OR = 1.49, 95% CI 0.79–2.81, p = 0.24). Thus, although cancer samples exhibited broader mature-domain peptide diversity, quantitative PSM analyses revealed persistent and significantly enriched pro-domain representation for NGF.

The relative composition of pro- vs mature-forms widely varies across cancer tissues (Figure 5C). In particular, Primary Biliary Cholangiocarcinoma (PBC), Lung Squamous Cell Carcinoma (LSCC), and Cancer Cell Line Encyclopedia (CCLE) datasets only showed profiles corresponding to the *BDNF* mature form (Figure 5C). Additionally, Lung Adenocarcinoma (LUAD) and hepatocellular carcinoma (HCC) also showed a strong bias toward the mature domain in comparison to the propeptide region (Figure 5C), which is in line with findings that BDNF/NTRK2 signaling encourages invasion, metastatic spread, and anoikis resistance in epithelial cancers (23,24). On the other hand, in oral squamous cell carcinoma (OSCC), only peptides mapping to the proBDNF form are observed.

Peptides corresponding to the proNGF form are prevalently found in most cancer tissues, and in particular in CCRCC, LSCC, OSCC and PDAC (Figure 5C). Tissue-resolved Fisher’s exact tests corroborated this heterogeneity (Figure 5E): the global NGF pro-domain enrichment was driven predominantly by BRCA, COAD and GC (all p < 0.0001), whereas LUAD showed a significant, opposite shift toward mature-domain enrichment for both NGF (p = 0.0037) and BDNF (p = 0.0004), wherein the latter is the principal contributor to the otherwise non-significant global BDNF trend.

On the other hand, NGF exhibited substantial propeptide representation alongside mature fragments, suggesting that precursor and mature forms coexist across multiple cancer contexts.. Positional scoring (supplementary Figure 7), which shows that maximal identification peaks for both proteins were localized in the C-terminal of the mature sequences, especially in aggressive cancers like LUAD and PBC, further supported the dominance of mature fragments. The idea that the mature NGF signaling axis is hyperactivated, possibly enhancing tumor–nerve crosstalk and progression processes that are becoming more widely recognized as drivers of poor prognosis across solid malignancies, is supported by high identification scores for NGF in these aggressive cancers (25). Together, these results suggest that the predictive behavior of NGF and BDNF in various cancer types may be significantly influenced by domain-specific proteolytic maturation rather than total protein abundance alone.

### Alternative spliced forms of innervation genes are differentially expressed in cancer and connected to distinct molecular functions

We considered protein coding alternative spliceforms of innervation genes. We found a total of 82 spliceforms, corresponding to 20 unique genes, differentially expressed in at least one of 16 cancer tissues (Figure 6A; Supplementary Table 2; see Methods). The three genes with most significantly DE splicevariants are *GNAS*, *NTRK3* and *ARTN*. Deletions, identified with respect to the protein canonical isoforms, is the most frequent type of splice variations, while the most common functional consequences on protein sequence is at the level of PTM sites as well as on binding regions involved in protein-protein interactions (PPIs) (Figure 6A). We found 12 genes characterized by at least two splice variants engaging with distinct sets of interacting proteins as well as being connected to distinct biological processes (Supplementary Figure 8). Among them is *NTRK3*, which is characterized by multiple splice variants that are significantly DE in different cancer tissues (Figure 6B). For instance, the 2nd splice form of *NTRK3*, i.e. Q16288-2, is over-expressed in kidney and skin, while under-expressed in prostate, lung, thyroid and colon cancer tissues (Figure 6B). This isoform is characterized by a large truncation of the kinase domain (Figure 6B) as well as by a completely different interactome (Figure 6C, violet) with respect to the canonical isoform (Figure 6C, green). Indeed, certain interactors reported to bind to *NTRK3*’s kinase domain (e.g. *DOK6* and *AP2B1*; Figure 6D), interact only with *NTRK3*’s canonical isoform and not with the 2nd isoform (Figure 6C). The different interactomes engaging with the canonical and 2nd isoform of NTRK3 are also connected to different biological processes (Figure 6E). Indeed, while the canonical isoform is associated with neurotrophin signaling pathways (e.g. “Brain derived neurotrophic factor BDNF signaling pathway”, FDR=9e-10), the interactome of the 2nd isoform is more significantly associated to pathways such as “PI3K-Akt signaling pathway” (FDR=0.00023) (Figure 6E).

**Figure 6.**
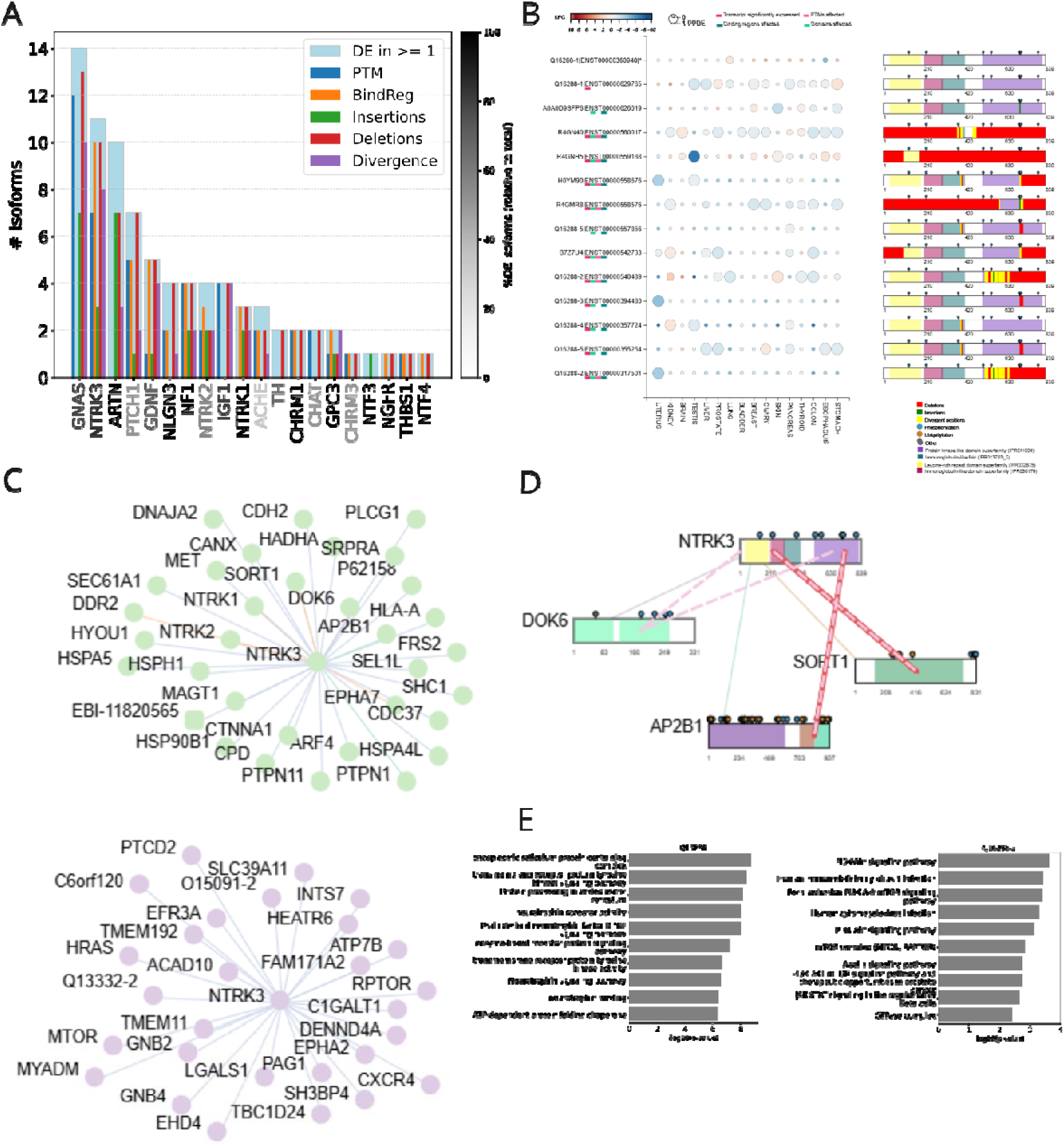
Differential expression and functional divergence of innervation gene spliceforms in cancer. **A**) The overlaid light-blue bars display frequency of protein-coding spliceforms of innervation genes, that are differentially expressed (DE) in atleast one cancer tissue. Within each blue bar, five stacked bars represent the number of DE splice-forms with different functional alterations. The gene labels on the x-axis are color-coded with a greyscale depicting the fraction of spliceforms relative to the total number of isoforms known for that gene. **B**) The bubbleplot on the left depicts the differential expression patterns of NTRK3 spliceforms across cancer types, wherein the color scale corresponds to the LFC values and bubble sizes are proportional to PPDE values. The cartoon panels to the right provide graphical representations of splicing-related variations, affected domains, and PTMs on the canonical protein sequence for NTRK3 isoforms. An adjacent legend clarifies the color coding for each feature. It is observed that the spliceform Q16288-2 shows opposing expression trends in different tissues and a truncated kinase domain. **C**) Distinct interactomes of NTRK3 canonical (green) and Q16288-2 (violet) spliceforms, indicating altered protein-protein interactions. **D**) Select interactors that bind to the kinase domain of canonical NTRK3 but are absent in the interactome of Q16288-2. The edge-colors represent different types of interactions while the solid red edges show the altered binding regions. This information is presented in the adjacent legend. **E**) Functional enrichment of interactomes, showing the canonical NTRK3 isoform is associated with certain processes (for eg. neurotrophin signaling), whereas Q16288-2 is linked to different distinct processes such as PI3K-Akt signaling, suggesting pathway divergence.

### Somatic alterations and expression of innervation genes are associated with patient survival

We considered somatic alterations and expression signatures of innervation genes to evaluate their capability to stratify patients based on their survival, whose data were retrieved and merged from three different sources (i.e. TCGA, ICGC and cBioportal; see Methods). When considering patients with mutations on any of the 26 innervation genes, we found significantly lower survival compared to patients with no alterations on these genes at the pancancer level (Figure 7A, Supplementary Figure 9; Logrank P-value=3.6e-5). Overall, we found that the majority of genes significantly associated with survival (9 out of 12; Logrank P-value < 0.05), are associated with lower prognosis (Figure 7A). The somatic alterations most significantly associated to lower survival across cancers are the ones affecting Neurotrophin-3 *NTF3* (Figure 7A,B; Logrank P-value=1.13e-05) and Neuroligin-3 *NLGN3* (Figure 7A,C; Logrank P-value=1.3-05). Other genes most significantly associated with lower survival are *GPC3*, *NGF* and *TH* (Figure 7A,D). As expected, we found a significant association of *NLGN3* to lower survival in glioma, together with *ACHE* and *NTRK3* (Figure 7F). In melanoma we found a significant association to lower survival of patients with somatic alterations of *NGF* (Figure 7E,F). This finding correlates with the recent report of an immunosuppressive effect elicited by the NGF–TrkA axes in melanoma which hampers durable memory T cell protection (26). When performing the survival analysis on patients carrying somatic alterations at interacting gene pairs, we found that the stratification potential of *NGF* mutations for patients with lower survival increases when considered in combination with mutations at *NTF3* (P-value=8.52e-4) and *NGFR* (P-value=7.37e-4;Figure 7G). Also the Brain derived neurotrophic factor *BDNF* enhances its association with lower survival when its mutations are considered together with *ACHE* at a pancancer level(Figure 7G). In individual cancer tissues, we found that patients affected by mutations of the *GDNF-NGFR* axes were significantly associated with lower survival in hepatobiliary carcinoma tissues (Figure 7G).

**Figure 7:**
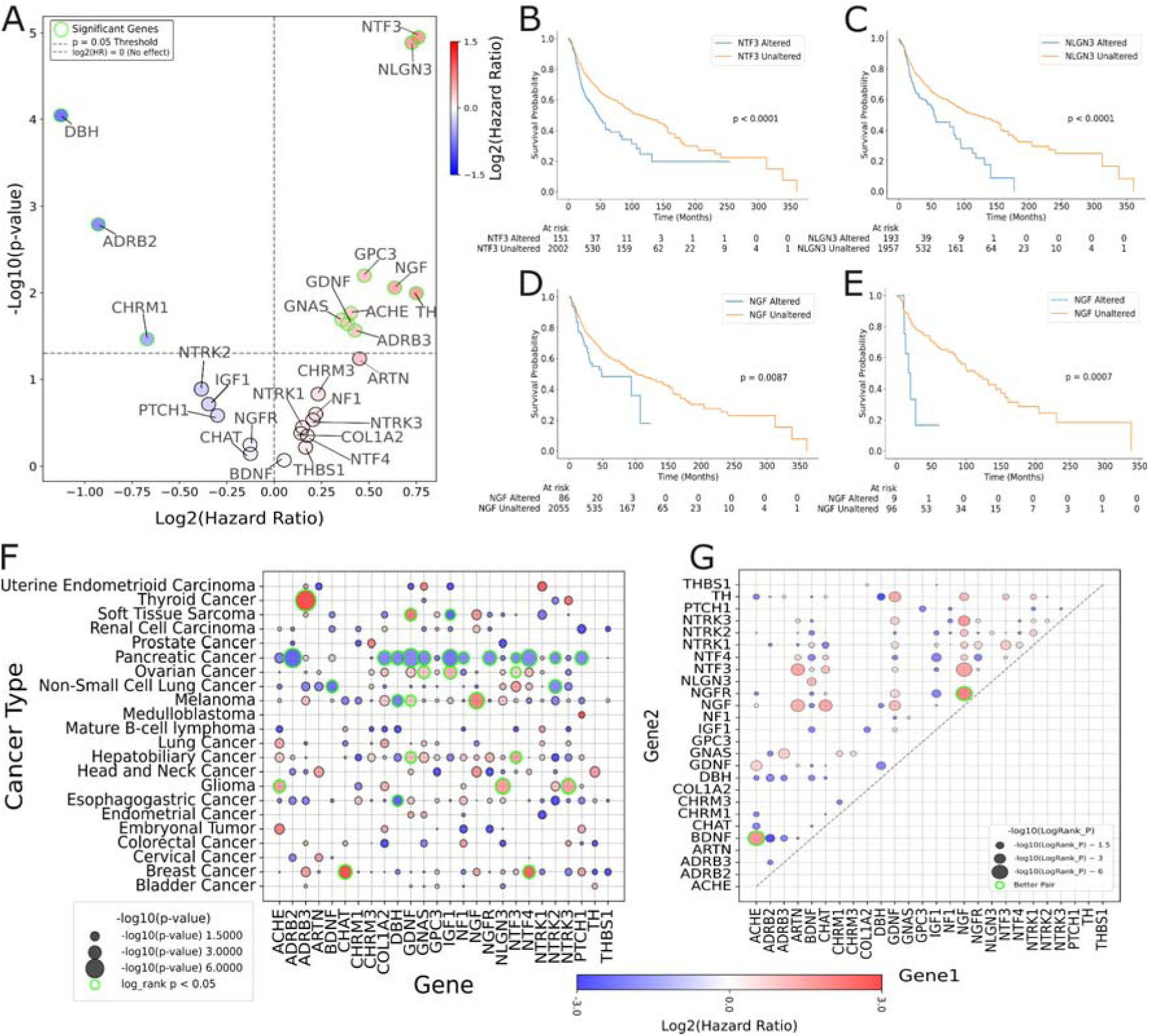
Survival analysis employing mutational data. A) The volcano plotdepicts −log10(p-value) and log2 (hazard ratio)of individual genes across Pan-Cancer. B) KM plot of NTF3 altered and unaltered in Pan-Cancer. C) NLGN3 KM plot in Pan-Cancer shows the altered and unaffected group. D) KM plot of NGF altered and unaltered in Pan-Cancer. E) KM plot of altered and unaltered NGF in melanoma cancer. F) A bubble map that depicts log-rank p value and hazard ratio individual genes across distinct cancer types. F) The bubble map depicts the logarithm base 10 of the p-value and logarithm base 2 of the hazard ratio of interacting gene pairs at the Pan-Cancer level. The green border color in A and B reflects the statistically significant genes with p < 0.05, while in F it signifies the more effective interaction gene pairs. In the bubble map, the patient size was set at ≥5. The size of the bubble indicates the logarithm base 10 of the p-value, while the hue of the bubble denotes the logarithm base 2 of the hazard ratio. Red color indicates lower survival and Blue color shows the higher survival.

*NGF,* together with *COL1A2* and *THBS1*, are also the single genes most significantly associated with lower survival when considering gene expression (bulk RNAseq) as a means to stratify patients (Figure 8A). In details, patients with higher expression of *NGF* have a significantly lower survival (P-value=3.26e-157). In general, there is a strong prevalence of significant associations to lower survival, with only five genes being associated with higher survival pancancer (Figure 8A). When considering functional interacting gene pairs, the widespread association to lower survival at the pancancer level is further reinforced (Figure 8B). We found several instances of gene pairs whose concordant expression in the same sample, either above or below the median, is associated with survival more significantly than by considering individual genes (see Methods). We found the strongest synergistic association in stratifying patients at higher risk at a pancancer level, when considering NGF in combination with the Tyroxine Hydroxylase (TH), a key enzyme in the biosynthesis of catecholamines (P-value=2.84e-17, Figure 8B,C). We also found that considering the combined expression of *NGF* with either *NGFR* or *BDNF* allowed us to stratify patients at higher risk compared to the individual instances (P-value=1.2e-157 and P-value=8.44e-232, respectively). Patient stratification also improves when considering *BDNF* in combination with *NTF4* and *TH*. At the individual tissue level, we also found multiple significant associations to cancer such as stomach, kidney, lining of body cavities, eye, brain or bladder both at the individual genes,(Figure 8D), as well as the pair level (Supplementary Figure 10). In particular, in stomach cancer we identified many significant instances associated to lower survival involving components of the neurotrophin and acetylcholine signaling pathways (Figure 8D, Supplementary Figure 10), further highlighting the role of the cross-talk of these pathways in supporting gastric cancer tumorigenesis (11).

**Figure 8:**
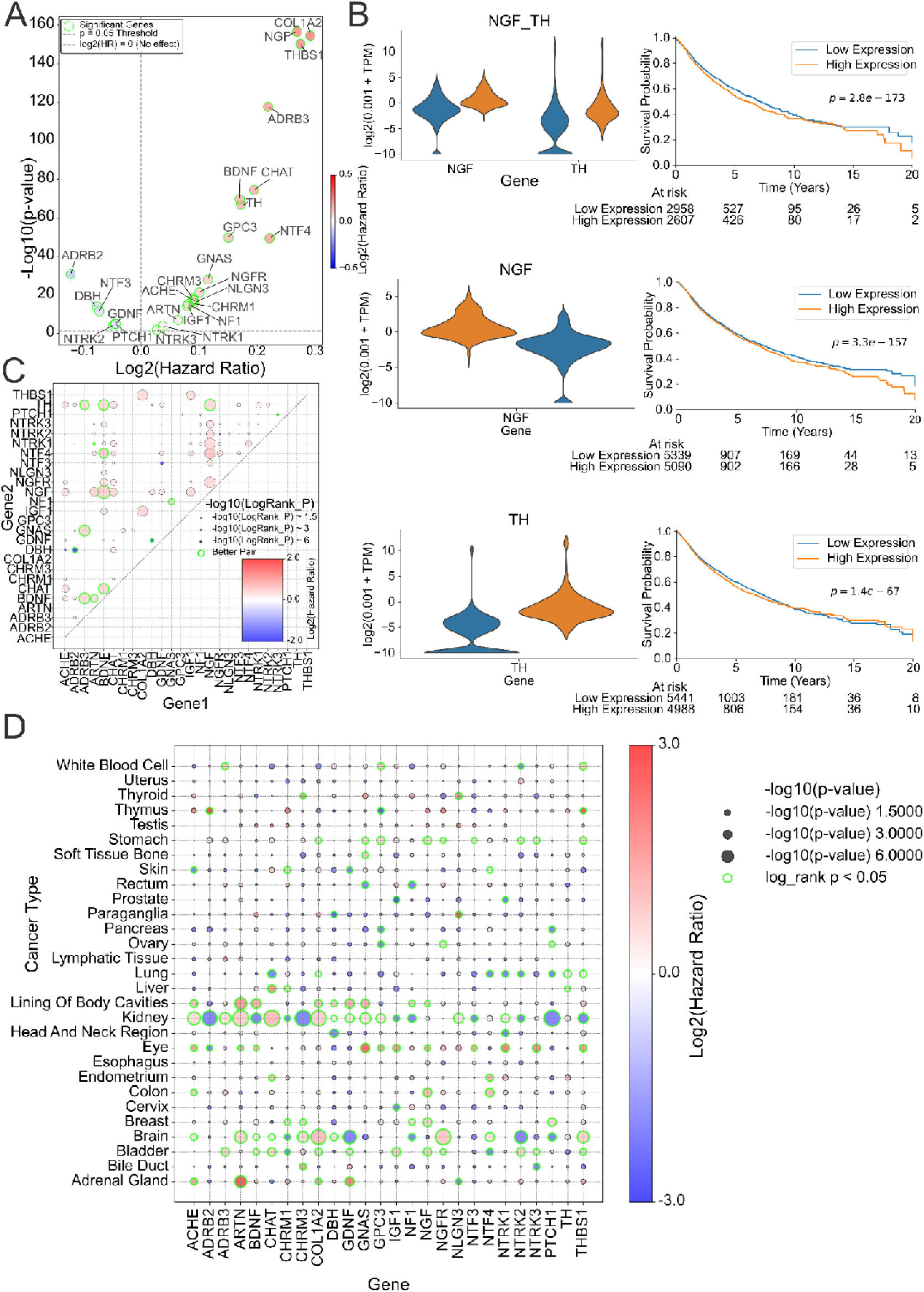
Survival analysis utilizing expression data from TCGA. A) The volcanoplot illustrates the −log10(p-value) and log2 (hazard ratio) of innervation pathway genes across Pan-Cancer. B) Violin plots illustrate the distribution of individual gene expression for NGF and TH, whereas KM plots show enhanced patient stratification based on the combined gene expression of NGF and TH in comparison to individual genes. C) The Bubble map exhibiting the −log10(p-value) and log2 (hazard ratio) of interacting gene pairs throughout the Pan-Cancer analysis. D) The bubble map depicts the −log10(p-value) and log2 (hazard ratio) of individual genes across different types of cancer. The green edge color in A and D represents the significant genes with p < 0.05, whereas in C it shows the more efficient interacting gene pairings. Examine cases where each stratified group comprises a minimum of five patients. The bubble’s size signifies the base −10 log of the p-value, whereas the bubble’s color represents the log2 (hazard ratio). Red color represents lower survival and blue color indicates higher survival.

## Discussion

In this study we have analyzed large scale cancer genomic datasets to identify omic signatures associated to genes involved in tumor innervation. We found a widespread mutational profile, with a prevalence of amplification events, which suggests that innervation genes might overall play an oncogenic role in cancer. Altogether, mutations of these genes affect 65% of patients in a pancancer cohort. Genes of the neurotrophin pathways are the most mutated ones, affecting 42% of patients pancancer. Such numbers further support the emerging role of neuronal factors and processes as modulators of cancer hallmarks, and overall contributing to tumorigenesis in multiple cancer contexts (1,2).

We found out that *NTRK1* is among the three most mutated innervation genes, mainly affected by amplifications. *NTRK3* appears as the most variable receptor when considering cancer somatic mutations as well as splice variants. Both genes, together with NTRK2, are reported to be cancer driver genes in both the Cancer gene census (27) as well as OncoKb (28) and to play an oncogenic role via fusion rearrangements. Neurotrophin ligand genes are not reported as oncodrivers. Consistently, we found that neurotrophins are affected by a lower number of mutations compared to cognate receptors. This pattern is similar to what has been found for other TRK systems, where the receptor is mutated, through point mutations, deletions or fusion, to destroy the receptor’s off-switch and keep the receptor’s signaling constitutively active. On the other hand, cancer cells do not need a mutated ligand to get continuous growth signals. Instead, they simply overexpress wild-type ligands, forcing their own receptors into overdrive(29). Perturbation of the neurotrophin genes appears to be highly detrimental, since both somatic mutations as well as higher expression of these genes are strongly associated with patients’ poor prognosis. In particular, we found that several neurotrophins are heavily associated with lower survival, either when considering mutations, e.g. affecting *NTF3* or *NGF*, as well as expression, e.g. of *NGF* and *BDNF*, in specific cancer tissues as well as across cancers. Considering combination of alterations for the genes reported to be in functional interaction further improve the stratification of the patients and identification of groups with worse prognosis, further suggesting that their interplay in mediating tumor innervation is associated with cancer severity. Indeed, we found several instances of receptor-ligand pairs (e.g. NGF-NGFR) significantly associated with lower prognosis, and better than the individual gene instances. We have also found several combinations of genes participating in different pathways that enhance patient stratification, such as TH with NGF, which highlights the role of the crosstalk between catecholamine and neurotrophin signaling in establishing a feed forward loop sustaining innervation and tumorigenesis. Many of the significant associations with worse prognosis have been also obtained at a pancancer level, as well as across multiple cancer types, suggesting that the crosstalk mechanisms reinforcing tumor innervation might indeed be a more widespread mechanism than previously thought.

In our analysis we also considered additional omics data, including alternative splicing isoforms, which revealed additional layers of regulation of neurotrophin and innervation pathway signaling, with alternative splicing architectures characterized by a substantial rewiring of their interaction networks. Likewise, analysis of neurotrophins proteoforms from proteogenomics datasets illuminated the specific regulation of the precursor as well as mature forms of neurotrophins in cancer. ProBDNF can signal through p75NTR complexes to induce apoptosis or growth suppression, while mature BDNF preferentially activates TrkB-mediated prosurvival pathways. This shift may help explain the positive survival associations found for BDNF in specific tumor contexts (22,30).

We integrated differential expression profiles from various cell types derived from tumour single-cell datasets to enhance these observations. The widespread dysregulation of NGFR from scRNAseq data aligns with previously reported studies on the role of NGFR in cell survival or apoptosis based on the cellular context (31). NGF overexpression was primarily detected in fibroblast populations, as opposed to malignant epithelial cells. This finding aligns with recent evidence suggesting that adrenergic stimulation of cancer-associated fibroblasts (CAFs) via ADRB2 signalling facilitates NGF secretion, thereby creating a feed-forward loop that increases tumour innervation and norepinephrine accumulation within the tumour microenvironment (32). These findings collectively support a model in which *NGF* functions as a paracrine mediator of tumor-stroma-nerve interactions, linking neural and microenvironmental signals to cancer development. Other neurotrophin axes contribute to the tumour–stromal interactions. *BDNF* and *NTRK2* were concurrently up-regulated in various endothelial and malignant cells, aligning with findings that tumour cells create autocrine loops which reinforce tumor survival and invasion through self=sustaining mechanisms (33,34). *NGF* and *NGFR* were similarly enriched in fibroblasts, mast cells, and stromal populations, consistent with studies demonstrating NGF/NGFR-mediated interactions between fibroblasts and mast cells that promote stromal activation and tumour progression (35). *NTRK1* showed up-regulation in fibroblasts and mast cells, while down-regulation occurred in monocytes/macrophages. In contrast, *NGF* remained up-regulated in immune cell types. This pattern indicates that *NGF* may sustain paracrine signalling and affect immune function despite low levels of its cognate receptor, aligning with emerging evidence that *NTRK1* and *NGF* signalling regulate macrophage activity and immune checkpoint responses (36,37). *NTF3* and *NTRK3* showed marked up-regulation in endothelial cells, remarking the role of *NT3–NTRK3* signalling in vascular remodelling and angiogenesis (38).

Moreover, while TRK receptors frequently undergo mutations or fusions leading to ligand-independent activation, our findings indicate that neurotrophins ligands are predominant overexpressed in stromal and immune cells, and their over-expression is significantly associated with lower survival of cancer patients. Overexpression enhances paracrine signalling that affects angiogenesis, immunological regulation, and tumour innervation.

Further analyses will be required to better understand the role of these genes in tumor innervation and their cross-talk with other signaling pathways. Analysis of scRNAseq and spatial transcriptomics data will be instrumental in revealing a more mechanistic picture of the interplay of these genes in mediating tumor innervation and will offer new precision, synergistic therapeutic opportunities.

## Methods

### Study Design and Gene Set Definition

To explore the neuronal genes’ involvement in cancer innervation pathways, we selected 26 genes on the basis of their reported role in neurotrophins pathway (*NGF, NGFR, BDNF, IGF1, GDNF, ARTN, NTF3, NTF4, NTRK1, NTRK2, NTRK3*), norepinephrine signaling pathway (*ADRB2, ADRB3, GNAS, DBH, TH*), acetylcholine signaling pathway (*ACHE, CHRM1, CHRM3, CHAT*), paracrine neuronal factors (*COL1A2, NLGN3, NF1*) and synaptogenic factors (*GPC3, PTCH1, THBS1*) (1,2,39–41).

Interactive gene pairs were obtained for the above selected genes using the STRING database (42), a database of experimental and predicted protein–protein interactions. We applied the default STRING settings, including a confidence score threshold of 0.04 for interactions supported by co-expression, experimental evidence, text mining, and related evidence channels. This resulted in a network consisting of 109 gene pairs with functional associations that were used in downstream analyses.

### Genomic Alteration Data and Clinical Cohort Assembly

For the pan-cancer study, we downloaded the *Pan-cancer analysis of whole genomes (ICGC/TCGA, Nature 2020)* dataset from cBioPortal (43).

https://www.cbioportal.org/results/mutualExclusivity?cancer_study_list=pancan_pcawg_2020&Z_SCORE_THRESHOLD=2.0&RPPA_SCORE_THRESHOLD=2.0&profileFilter=mutations%2Ccna%2Cmrna_seq_fpkm_capture_all_sample_Zscores&case_set_id=pancan_pcawg_2020_all&gene_list=NGFR%250ANTF3%250ANTF4%2520%250ABDNF%250ANGF%250ANTRK1%250ANTRK2%250ANTRK3%250AIGF1%250AGDNF%250AARTN%250AACHE%250ACHRM1%250ACHRM3%250ACHAT%250AADRB2%2520%250AADRB3%250ATH%250ADBH%250AGNAS%250AGPC3%250ATHBS1%250APTCH1%250ANLGN3%250ANF1%250ACOL1A2&geneset_list=%20&tab_index=tab_visualize&Action=Submit

Since cBioPortal only included a selected set of clinical studies, we downloaded additional TCGA and ICGC data (January 2024 release) to maximize clinical coverage. TCGA data were retrieved from the Genomic Data Commons:

https://portal.gdc.cancer.gov/analysis_page?app=

Specifically, the file *Clinical with Follow-up-clinical_PANCAN_patient_with_followup* was utilized, and the column *bcr_patient_barcode* was used as a key for merging. ICGC data were downloaded from the International Cancer Genome Consortium (www.icgc.org), and the column *icgc_donor_id* from the relevant file was used for the merge operation.

Data preprocessing and merging were performed in R (v4.4.3). The cBioPortal dataset was first merged with the ICGC dataset and subsequently with the TCGA dataset using the identifiers *icgc_donor_id* and *bcr_patient_barcode*, respectively. After merging, overall survival time was computed and patients without survival information were excluded. The final extended dataset contained 2,214 patients out of a potential 2,583.

Binary alteration matrices, alteration-type summaries, mutation annotations, OncoPrint data, and mutual exclusivity/co-occurrence statistics were downloaded from cBioPortal following gene-based queries. The downloaded file *“Sample matrix: List of all samples where 1 = altered and 0 = unaltered”* was used to determine alteration status for each gene across samples. Additional alteration-type information was obtained from the *“Type of Genetic Alterations Across All Samples”* table, while somatic mutation classifications were downloaded from the *Mutations* tab.

Alteration classes included mutations, copy-number alterations, and expression-based events. The resulting alteration matrix was merged with the extended clinical cohort to generate the mutational-survival dataset containing both survival information and alteration profiles.

### Gene Fusion Dataset

Gene fusion data were obtained from FusionGDB2 (https://compbio.uth.edu/FusionGDB2/download.html) (44). Specifically, the file *All gene symbols arranged fusion genes* was downloaded together with head and tail breakpoint annotations.

We refined the gene fusion dataset by retaining fusion events in which either the 5′ partner (Hgene) or the 3′ partner (Tgene) corresponded to one of the genes in our candidate gene set. Duplicate entries and non-cancer categories were removed. The number of unique sample IDs was calculated for each fusion event across cancer types. Visualizations were generated using matplotlib (v3.8.2).

Fusion breakpoint coordinates for *NTRK1, NTRK2*, and *NTRK3* were characterized as Head (5′ partner) or Tail (3′ partner retaining the tyrosine kinase domain) and organized according to genomic position to determine breakpoint frequencies across samples.

Breakpoint coordinates were reported relative to GRCh37/hg19 and converted to GRCh38/hg38 using the UCSC LiftOver tool implemented through rtracklayer (v1.64), restricting mappings to ±50 kb around each gene locus to eliminate false-positive mappings.

Annotations for genes, transcripts, and Pfam protein domains were obtained from Ensembl GRCh38 release 109 using the AnnotationHub and ensembldb Bioconductor packages. Canonical coding transcripts were selected for each gene. Breakpoint distributions were visualized relative to transcript structures and kinase-domain positions using Gviz (v1.48). All analyses were performed using R (v4.4.3; Bioconductor v3.20).

### Bulk Transcriptomic Datasets

Expression data from a combined TCGA, TARGET, and GTEx cohort (45) were retrieved from the UCSC Xena Browser (46) using the UCSC Toil RNA-seq Recompute resource. Only TCGA primary tumor samples and GTEx normal tissues were used in downstream analyses. Both raw count matrices and log-transformed TPM matrices [log2(TPM + 0.001)] were analyzed.

For co-expression analyses, expression data were partitioned into tissue groups according to UCSC Xena metadata. TCGA and GTEx cohorts were processed separately. Count matrices were normalized using PyDESeq2 (v0.4.4), followed by variance-stabilizing transformation.

### Single-Cell Transcriptomic Datasets

Pre-calculated differential expression data for neurotrophin genes BDNF, NGF, NGFR, NTRK1–3, GDNF, ARTN, NTF3, and NTF4 were obtained from the TISCH single-cell RNA-seq database (18). In TISCH, differential expression is calculated by comparing each annotated cell type against all other cell types within the same dataset.

Genes with adjusted P values < 0.05 were considered significant. Duplicate entries sharing identical tissue, cell type, dataset, and log fold change values were removed while retaining the most significant observation. Results were aggregated at both tissue and cell-type levels. Cell annotations were consolidated into major categories including Fibroblast, Malignant, T/NK, Mono/Macro, B, DC, Endothelial, Epithelial, Neural, and Others. Gene regulation direction was determined according to the sign of log2 fold change.

### Alternative Splicing Resources

We used the EXPANSION API (https://expansion.bioinfolab.sns.it/api/) to collect alternative spliceform-specific information for our candidate genes [cite expansion]. Retrieved information included differentially expressed alternative spliceforms in cancer (TCGA) versus normal tissues (GTEx), splicing events, spliceform-specific interaction networks, over-representation analyses, and spliceform-specific functional effects such as altered PTM sites and binding regions.

For data retrieval and parsing, an in-house Python (v3.9.7) script was used to perform HTTP POST requests using requests (v2.31.0). API responses were parsed using the built-in json package. Downstream processing was performed using pandas (v2.3.0), while visualization was generated using matplotlib (v3.8.2) and seaborn (v0.13.1).

### Proteomic Analysis

To verify the existence of specific NGF and BDNF peptide sequences across 48 proteomic datasets in the PepQuery library (47), peptide-centric identification was performed using the standalone version of PepQuery (v2.0.2).

Target peptide lists were generated through in silico digestion of human NGF and BDNF protein sequences, allowing one missed cleavage, as identified using MS-Digest from the University of California, San Francisco.

To accommodate different labeling chemistries, separate modification profiles were used. For TMT-labeled datasets, oxidation (M) and phosphorylation (S, T, Y) were specified as variable modifications, while carbamidomethylation (C) and TMT 11-plex (K, N-term) were specified as fixed modifications. For iTRAQ-labeled datasets, iTRAQ 4-plex (K, N-term) was specified as a fixed modification, whereas iTRAQ 4-plex (Y) was included as an additional variable modification.

PepQuery validation consisted of three sequential steps: (1) scoring target peptides against experimental spectra using the Hyperscore algorithm; (2) calculation of p-values using a decoy search against 10,000 random peptides; and (3) unrestricted PTM searching to determine whether the queried peptide remained the best explanation for the observed spectrum.

Only peptides classified as *Confident* by the PTM competition filter and associated with p < 0.05 were retained.

### Mutational Landscape Analysis

The genomic alteration landscape of the 26 candidate genes was assessed using OncoPrint summaries in cBioPortal. Alteration frequency was defined as the number of unique patients carrying an alteration divided by the total cohort size. Visualization was performed using Python v3.10.13 with pandas (v2.3.0), NumPy (v2.2.6), matplotlib (v3.8.2), and seaborn (v0.13.1).

Alteration frequencies by cancer type were obtained directly from cBioPortal. Oncogenic mutation classes (Missense, Truncating, In-frame indel, and Splice-site) were visualized as stacked bar plots. Lollipop plots for NF1, COL1A2, and NTRK3 were obtained directly from the Mutations tab of cBioPortal.

### Bulk Differential Expression Analysis

Differential expression analysis was performed following the methodology described in [REF CELL GENOMICS] using TCGA and GTEx count data retrieved from UCSC Xena. Candidate genes were classified as differentially expressed when |log2 fold change| > 1 and adjusted P < 0.05.

Differential expression analyses were performed using DESeq2 in R by contrasting TCGA tumor samples against GTEx normal tissues across 16 tissues. Visualization was performed using matplotlib (v3.8.2) and seaborn (v0.13.1).

### Single-Cell Differential Expression and Cell-Type Aggregation

Neurotrophin genes exhibiting adjusted P values < 0.05 were considered significant. Aggregated analyses quantified the number of datasets showing up- or down-regulation and computed mean log2 fold changes separately for up-regulated and downregulated observations. Visualization was performed using pandas (v2.3.0), NumPy (v2.2.6), and matplotlib (v3.8.2).

### Survival Analysis

#### Survival Analysis of Genomic Alterations

Survival analyses were performed using Python v3.10.13 and lifelines (v0.28.0). Two conditions were evaluated: (i) altered versus unaltered for individual genes and (ii) altered versus unaltered for gene pairs. Unaltered samples were used as the reference group. Overlapping patients were excluded from comparison groups.

Univariate Cox proportional hazards models were used to estimate hazard ratios and confidence intervals. Kaplan–Meier curves were generated using KaplanMeierFitter. Statistical significance was evaluated using the log-rank test, with p < 0.05 considered significant. Analyses were restricted to comparisons containing at least five patients per group.

Gene pairs were considered improved predictors when they satisfied all of the following criteria: concordance index ≥ 0.5, adjusted p-value < 0.05, and a log-rank p-value lower than that obtained for either constituent gene alone.

#### Survival Analysis of Gene Expression

Log-transformed TPM matrices from TCGA were used for expression-based survival analyses. Cox proportional hazards models were implemented using the R package survival (v3.7).

Each model was trained on patients belonging to a single tissue. Tumor types within the same tissue category were treated as separate strata. For individual genes, patients were dichotomized according to median expression. For gene pairs, patients were classified according to whether both genes were simultaneously above or below their respective median values. Samples with discordant expression states were excluded.

Kaplan–Meier curves were visualized using lifelines. Gene pairs were considered superior stratifiers only when Benjamini–Hochberg adjusted p-values were < 0.05 and all three metrics improved relative to both constituent genes: absolute Cox coefficient, concordance index, and log-rank significance.

Heatmaps were generated using matplotlib (v3.8.2) and seaborn (v0.13.1) with Ward hierarchical clustering. Statistical significance was defined as p < 0.05.

### Statistical Analysis

Data processing, statistical modeling, and visualization were performed using Python (v3.9.7 and v3.10.13) and R (v4.4.2 and v4.4.3; Bioconductor v3.20), employing pandas (v2.3.0), NumPy (v2.2.6), matplotlib (v3.8.2), scipy.stats (v1.12.0), and seaborn (v0.13.1).

A significance threshold of 0.05 was applied throughout. Benjamini–Hochberg false-discovery-rate correction (Padj < 0.05) was used for high-throughput analyses, including bulk differential expression, single-cell differential expression, and expression-based survival analyses. Unadjusted p-values (p < 0.05) were applied to low-dimensional hypothesis testing, including PepQuery validation and Kaplan–Meier survival comparisons, and Fisher’s exact tests comparing pro- versus mature-domain peptide-spectrum match (PSM) counts between healthy and cancer samples, performed both globally and per cancer type (scipy.stats.fisher_exact); odds ratios and 95% confidence intervals were computed on the log-odds scale with a Haldane–Anscombe continuity correction

Analyses required at least five patients per comparison group. Significant associations were defined using Cox model statistics, concordance index, and log-rank testing.

## Acknowledgments

F.R. was supported by the Italian Ministry of University and Research through the Department of Excellence “Faculty of Sciences” of Scuola Normale Superiore. The research leading to these results also received funding from the Italian Association for Cancer Research (AIRC) under My First AIRC Grant (MFAG) 2020 - ID. 24317 and Investigator Grant (IG) 2025 - ID. 32170 projects. F.R. was also supported by the Project granted by Next Generation EU – National Recovery and Resilience Plan (Piano Nazionale di Ripresa e Resilienza, NRRP) – Mission 4 Component 2 Investment 1.4 – Ministry of University and Research (MUR) Call N. 3277 Project Code ECS_00000017 MUR Directoral Decree n.1055, 23 June 2022, CUP B83C22003930001, project title “Tuscany Health Ecosystem – THE”, Spoke 8. F.R. also received funding from the ImmunoHub project (“Immunoterapia: cura e prevenzione di malattie infettive e tumorali (Immuno-HUB)”).

